# Salt Induced Transitions in the Conformational Ensembles of Intrinsically Disordered Proteins

**DOI:** 10.1101/2022.02.16.480648

**Authors:** Hiranmay Maity, Lipika Baidya, Govardhan Reddy

## Abstract

Salts modulate the behavior of intrinsically disordered proteins (IDPs). In low ionic strength solutions, IDP conformations are primarily perturbed by the screening of electrostatic interactions, independent of the identity of the salt. In this regime, insight into the IDP behavior can be obtained using the theory for salt-induced transitions in charged polymers. However, in high ionic strength solutions, salt-specific interactions with the charged and uncharged residues, known as the Hofmeister effect, influence IDP behavior. There is a lack of reliable theoretical models in high salt concentration regimes to predict the salt effect on IDPs. Using a coarse-grained simulation model for the IDPs and experimentally measured water to salt solution transfer free-energies of various chemical groups, we studied the salt-specific transitions induced in the IDPs conformational ensemble. We probed the effect of three different salts, ranging from protective osmolyte to denaturant, on five IDPs belonging to various polymer classes classified based on charge content. The transitions observed in the IDP conformational ensembles are dependent on the salt used and the IDP polymer class. An important implication of these results is that a suitable salt can be identified to induce condensation of an IDP through liquid–liquid phase separation.

## Introduction

Salts have significant impact on the conformations sampled by intrinsically disordered proteins/regions (IDPs/IDRs). As a result, salts can affect diverse cellular functions of IDPs^1–6^ such as cell signaling,^5,7^ stress granule assembly,^8,9^ heterochromatin formation,^10^ transcription, etc. Furthermore, transitions in the conformational ensembles sampled by IDPs due to the changes in cellular environment can result in pathologies^9,11^ such as neurodegenerative diseases^12^ and cancer^13^ through the formation of membraneless organelles^14^ via liquid–liquid phase separation (LLPS).^15–19^

Salts modulate a range of biophysical processes such as protein folding, aggregation, protein–RNA interactions, protein crystallization, and precipitation.^20–23^ The mechanism through which salts interact with biomolecules depends on the salt concentration ([*salt*]). In low [*salt*], the ions screen the electrostatic interactions between the charged residues independent of the salt used. However, in high [*salt*], the ions interact with the biomolecules through salt-specific interactions known as Hofmeister effects. Salt ions are arranged in a series referred to as Hofmeister series based on their ability to stabilize/destabilize (salting-out/in) the folded state of a globular protein.^23–25^

At physiological conditions, IDPs rapidly inter-convert between non-specific conformations as they are devoid of secondary/tertiary structures due to a significant fraction of polar and charged residues present in their sequences.^2,26,27^ There has been extensive effort using experiments and theory to understand the effect of salts on the IDPs.^28–31^ Single-molecule Forster energy transfer (smFRET) and simulations demonstrated that in low ionic strength solutions, the dimensions of the IDPs increased with the net charge.^28,29,32^ Polyampholyte-like IDPs exhibited compaction in dimensions due to interaction between opposite charges.^32–36^ Experiments further demonstrated using the polymer scaling laws that solvent quality of water to the IDPs depends on the IDP sequence composition.^30,37,38^ However, it is challenging to characterize the conformational ensembles sampled by IDPs as they are highly influenced by the charge distribution in the sequence. The charge distribution in the IDPs can be characterized by parameters such as the fraction of charged residues (*FCR*), the net charge per residue (*NCPR*), charge asymmetry (*σ*_±_), charge pattern (*κ*) and sequence charge decoration (*SCD*)^33,35,36,39–41^ (see Methods for details). These parameters further help identify the IDPs to the polymer class they belong to, such as polyelectrolytes, polyampholytes, or uncharged polymers.

Since charge distribution in the IDPs plays a critical role in influencing their conformational ensemble, salts will significantly impact IDP conformations and their physical properties.^42^ In low [*salt*] solutions, salts affect the IDP conformations through charge screening, independent of the salt identity. Therefore, we can exploit the existing polyelectrolyte^43–46^ and polyampholyte^34,39,47,48^ theories to understand the IDP behavior as demonstrated previously.^28,34,41^ However, in large [*salt*], salt identity is crucial due to the salt-specific Hofmeister effects^23,25^ that affect the IDP conformations. There is no analytic theory to account for the Hofmeister effects on IDPs. Recent coarse-grained hydrophobicity scale (HPS) model^31^ parameterized using the FRET efficiency experimental data was able to predict the salt effect on the LLPS of IDPs.

Computer simulations complement experiments and theory, and are playing an important role in elucidating the behavior of IDPs. Simulations were pivotal in understanding the IDPs conformational ensembles and dynamics.^32–34,49–57^ Computer simulations using atomistic force fields matching the appropriate length and timescales of the biophysical phenomena have the advantage of providing detailed information about the biophysical processes.^58–61^ However, the main drawbacks are the lack of reliable all-atom force fields to simulate the Hofmeister effect of salts, and it is computationally intensive to simulate large IDPs to obtain conformations representative of the equilibrium ensembles.

In this paper, we developed an efficient simulation model to compute the properties of IDPs in different salt concentration regimes and studied salt-induced transitions in the conformational ensembles of different classes of IDPs. To overcome the time scale problem associated with the sampling of representative IDP conformations, we used a coarse-grained model for the IDPs, which is a variant of the self-organized polymer model for IDP (SOP-IDP).^49^ The effect of salt on the IDP residues is taken into account using the molecular transfer model.^62,63^ The transfer energies of amino acids in various salt solutions are obtained from the experiments,^25,64^ where the salt’s effect on the solubility of model compounds used to mimic amino acids is measured and explained in terms of the solute partitioning model.^65,66^ We have successfully used this model in a previous study to probe the effect of salts on protein folding thermodynamics.^67^

To understand salt-induced transitions in the conformational ensembles of IDPs, we studied the effect of salt on the conformations of five different IDPs: (1) Nucleoporin, (2) Human prothymosin-*α*, (3) cyclin-dependent inhibitor kinase Sic1, (4) N-terminal transactivation domain (TADn) of ERM protein (ERMTADn), and (5) N-terminal domain of HIV-1 integrase (IN). These IDPs have different chain lengths (*N*_res_), compositions, sequences, and physicochemical properties based on the electrolytic behavior (Table S1). These IDPs are also used as model systems in experiments^28,68–71^ to understand IDP properties. Nucleoporin is an 81 residue long uncharged IDP rich in hydrophilic residues. The other IDP sequences contain ionizable charged residues and behave as polyelectrolytes or polyampholytes, depending on their *NCPR* and *FCR*. Human Prothymosin-*α* and Sic1 are polyelectrolytes, whereas ERMTADn and IN are polyampholytes. In the absence of Zn^2+^, IN is unstructured, and in this study, we treat it like an IDP. The IDP sequences are characterized^33,72,73^ using *FCR, NCPR*, and *σ*_±_ (Table S1).

To probe the effect of salts on IDPs, we performed simulations of IDPs in three different salt solutions: guanidine hydrochloride (GuHCl), potassium chloride (KCl), and potassium glutamate (KGlu). The salt GuHCl acts as a denaturant, which destabilizes a globular protein’s folded state, whereas KCl and KGlu are protective osmolytes, which stabilize the folded state. Experiments^28,30^ on IDPs show that GuHCl stabilizes the expanded conformations, whereas KCl and KGlu stabilize the compact conformations.

## Results and Discussions

### Kratky Plot of IDPs

We computed normalized intensity of scattered wave vector (*I*(*q*)/*I*(0)) (Eq. 2), and Kratky plot (*q*^2^*I*(*q*)/*I*(0)) for all the five IDPs using the simulation data at *T* = 300 K and [*salt*] = 0.15 M. The computed *I*(*q*)/*I*(0) and *q*^2^*I*(*q*)/*I*(0) are in near quantitative agreement with the measurements from small-angle X-ray scattering^68–71^ (SAXS) experiments in similar salt concentrations (Fig. 1A,B and S1). The agreement between simulations and experiments for both smaller (*q* ≲ 1.0 nm^−1^) and larger *q* values (*q* ≳ 1.0 nm^−1^) shows that the SOP-IDP model can capture the IDPs overall dimensions and accurately describe its structure at smaller length scales. The computed average radius of gyration 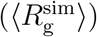 and the experimentally measured values 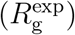 of IDPs are in good agreement (Fig. 1C). The Kratky plot of globular proteins exhibits a bell shaped curve due to the secondary and tertiary structure present in the protein. The Kratky plot of the IDPs exhibit a plateau at intermediate *q* values and further increases at larger *q* values indicating the absence of ordered structure (Fig. 1A,B).

**Figure 1:**
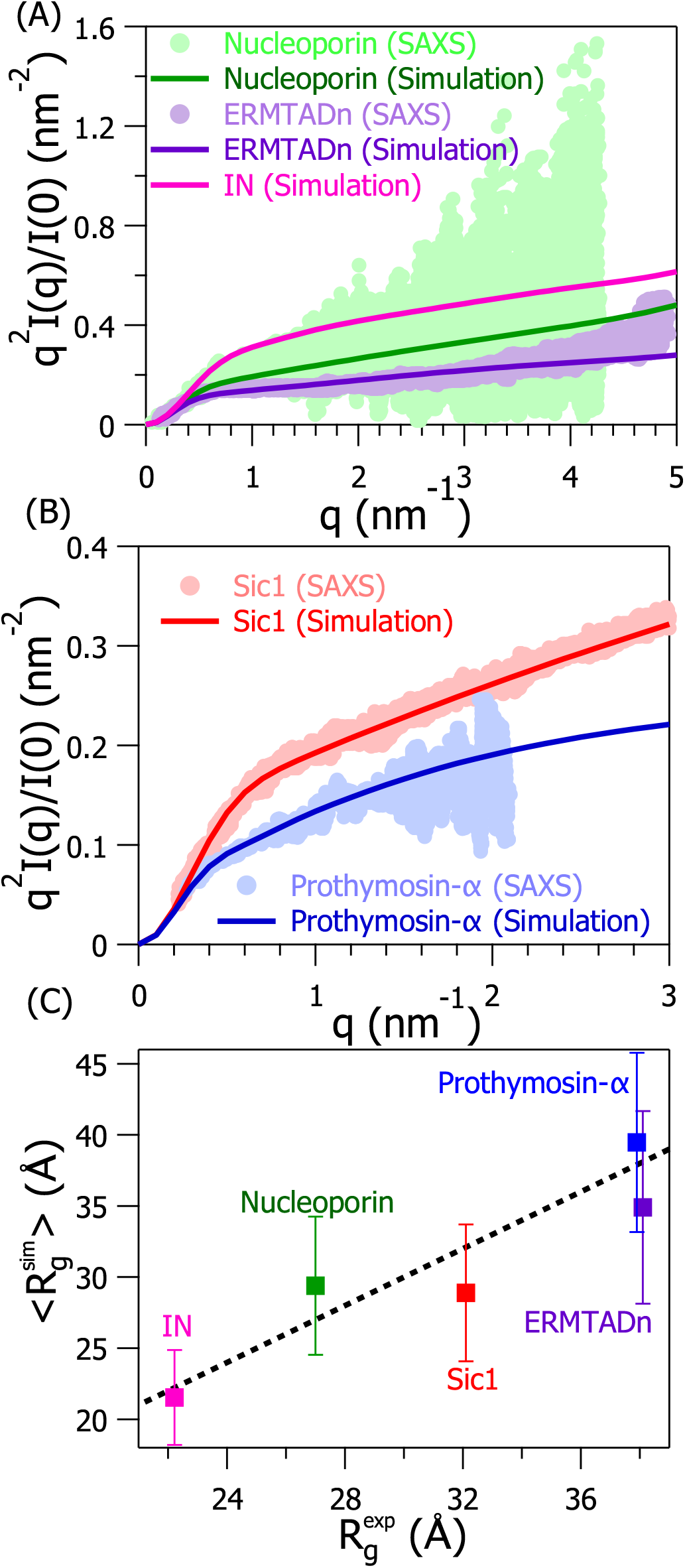
Normalized Kratky plot (*q*^2^*I*(*q*)/*I*(0) vs *q*) from experiments and simulations for (A) Nucleoporin (green), ERMTADn (violet) and IN (magenta), (B) Sic1 (red) and Prothymosin-*α* (blue). Experimental data is not available for IN. Light shaded dots and deep solid line are the corresponding experimental and simulated results, respectively. (C) IDPs average radius of gyration computed from simulation, 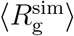, are plotted against the experimentally measured values, 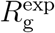. The Pearson correlation coefficient is ≈ 0.93.

### IDPs Exhibit Self Avoiding Walk (SAW) Behavior and Conformational Heterogeneity at Physiological Salt Concentration

The scaling exponent *ν*, which indicates the polymer behavior in a solvent is extracted using the scaling relation, *R*|*_i–j_*| ~ |*i* – *j*|*^ν^*, where *R*|*_i–j_*| is the distance between the residues *i* and *j* in the IDP. The exponent *ν* is also extracted from the IDP backbone structure factor (Eq. 3) using the scaling relation, *S*(*q*) ~ *q*^−1/*ν*^.^74^ The value of *ν* obtained from both the methods is ≈ 0.59 suggesting that IDPs behave like a polymer in a good solvent exhibiting the features of a SAW at physiological salt concentration (≈ 150 mM)(Fig. 2A,B).^34^ IDPs are heteropolymers and their conformational ensembles are highly heterogeneous in nature,^49,51,75–78^ which cannot be completely captured by the scaling exponents alone. We demonstrated the conformational heterogeneity in the IDPs by comparing their probability distribution of dimensionless end-to-end distance 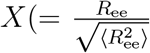, where *R*_ee_ is the end-to-end distance and 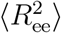 is the second moment of end-to-end distance) with the theoretically obtained universal expression for SAW^79–82^ (Fig. 2C). The universal shape of *P*(*X*) for the SAW is given by

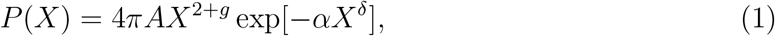

where *ν* is the Flory scaling exponent, *g* = (*γ* – 1)/*ν, δ* = 1/(1 – *ν*), and *γ* ≈ 1.1619 for 3 dimensional SAW.^83^ The constants, *A* and *α*, are obtained using the constraints 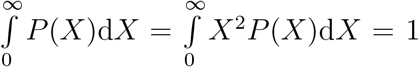. The simulated *P*(*X*) deviates significantly from the universal distribution, which can be attributed to the residue specific interactions and long range electrostatic interactions between the charged residues. In addition to *P*(*X*), the nonlocal contact (contact between two residues separated by ≥ 8 residues along the polypeptide chain) frequency obtained for a pair of residues is inhomogeneous unlike a homopolymer, which further indicates that IDPs sample a heterogeneous ensemble (Fig. S2).

**Figure 2:**
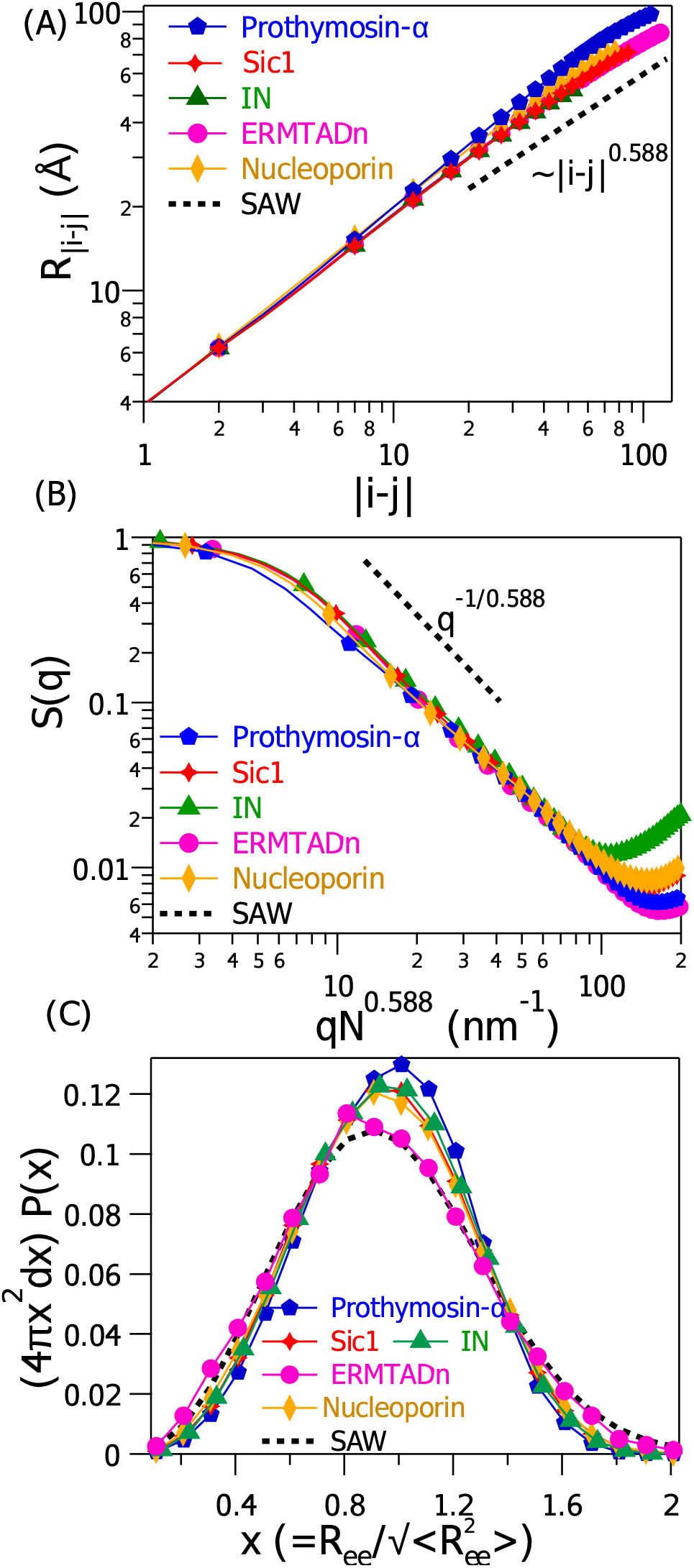
(A) Inter-residue distance (*R*|*_i–j_*|) is plotted as a function of separation between residues *i* and *j* (|*i* – *j*|) on a log-log scale for prothymosin-*α* (blue pentagons), sic1 (red diamonds), IN (green triangles), ERMTADn (magenta circles) and nucleoporin (yellow rhombus). The dotted line in black corresponds to a polymer exhibiting SAW for which *R*|*_i–j_*| ~ |*i* – *j*|^0.588^. (B) Structure factor (*S*(*q*)) for all the five IDPs is plotted as a function of scaled wave vector (*qN*^0.588^), where *N* is the number of residues in the IDP. The dotted line in black corresponds to a polymer behaving as a SAW for which *S*(*q*) ~ *q*^−1/0.588^. (C) Probability distribution of 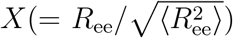, *P*(*X*) are plotted for all the five IDPs. The universal *P*(*X*) for SAW (Eq. 1) is shown in black dashed line.

### Electrostatic Interactions and Hofmeister Effects Modulate IDP Behavior in Low and High Salt Concentration Regimes

To probe salt induced transitions in the conformational ensembles of IDPs, we simulated IDPs in salt solutions of GuHCl, KCl, and KGlu. The salt GuHCl acts as a denaturant and destabilizes the folded state of globular proteins, whereas KCl and KGlu act as protective osmolytes, which stabilize the folded state.^28,30^ Salts modulate the IDPs behavior through the screening of electrostatic interactions and Hofmeister effects.^23,25^ In low salt concentrations ([*salt*] < 1.0 M), the effect of salt on the IDP is primarily through the screening of electrostatic interactions, and it is independent of the identity of the salt. However, in high salt concentrations ([*salt*] ≥ 1.0 M), the effect of salt on the IDP is through Hofmeister effects, and it is salt specific.^23,25^ To demonstrate this in the simulations, we performed different sets of simulations using the energy functions, Eq. S1 and S6. When we simulate the IDP using the energy function given by Eq. S1, we take into account only the screening of the electrostatic interactions in the IDP due to the addition of the salt. However, when we simulate the IDP using the energy function given by Eq. S6, we include the transfer energy (Δ*G_tr_*({**r**}, [*salt*])) in the energy function, which takes into account the salt specific Hofmeister effects on the IDP. Simulations show that the average radius of gyration 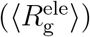 (Eq. 4) for a charged IDP initially increases or decreases due to charge screening for [*salt*] < 1 M when simulations are performed using Eq. S1 (Fig. S3). For [*salt*] ≥ 1 M, the effect of charge screening on 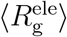 is minimal. However on adding the Δ*G_tr_*({**r**}, [*salt*]) term to the energy function (Eq. S6), the average radius of gyration (〈*R*_g_〉) of the IDPs increased in the case of GuHCl, which acts as a denaturant for folded proteins for [*GuHCl*] > 1 M due to salt specific Hofmeister effect (Fig. S3). This demonstrates that initially the effect of salt on the IDP is due to the salt identity independent screening of electrostatic interactions, and in high [*salt*] the effect on the IDP is due to the salt specific Hofmeister effect^23,25,64,84^ (Fig. 3, 4 and 5).

**Figure 3:**
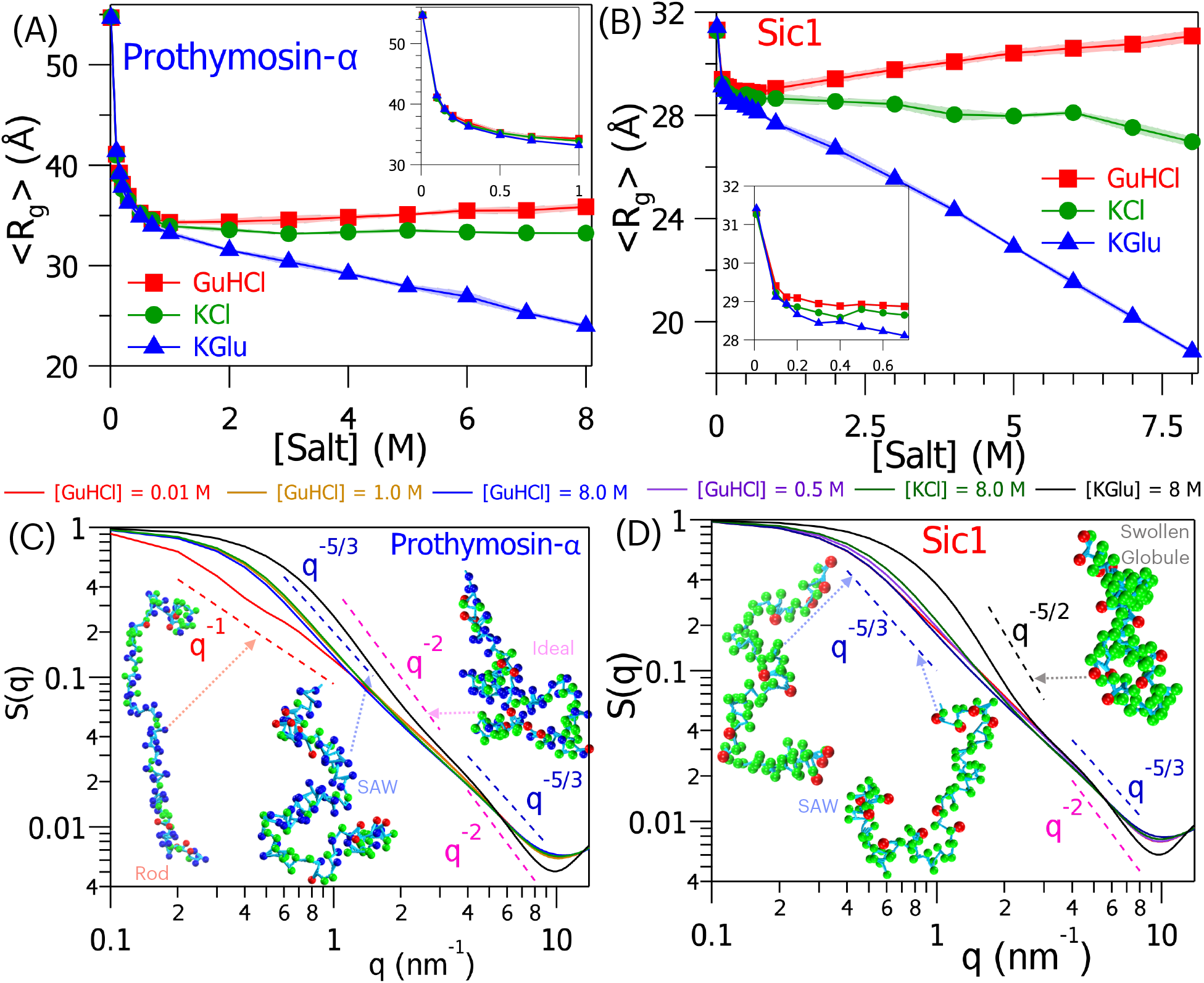
Average radius of gyration (〈*R*_g_〉) of (A) Prothymosin-*α* and (B) Sic1 computed as a function of [*GuHCl*] (red squares), [*KCl*] (green circles) and [*KGlu*] (blue triangles). 〈*R*_g_〉 as a function of [*salt*] in low [*salt*] regime (0 < [*salt*] ≤ 1 M) is shown in the inset. (C) Normalized structure factor *S*(*q*) of Prothymosin-*α* at [*GuHCl*] = 0.01 M (red), 1.0 M (yellow) and 8 M (blue), [*KCl*] = 8 M (green) and [*KGlu*] = 8.0 M (black). (D) *S*(*q*) for Sic1 at [*GuHCl*] = 0.01 M (red), 0.5 M (violet) and 8 M (blue), [*KCl*] = 8 M (green) and [*KGlu*] = 8.0 M (black). The dashed straight lines with scaling *q*^−1^, *q*^−5/3^, *q*^−5/2^, and, *q*^−2^ correspond to rod-like (red), SAW (blue), swollen globule (black), and, ideal (magenta) configuration. Representative IDP structure(s) are shown in bead representation. The acidic, basic and neutral residues are shown as blue, red and green beads, respectively.

**Figure 4:**
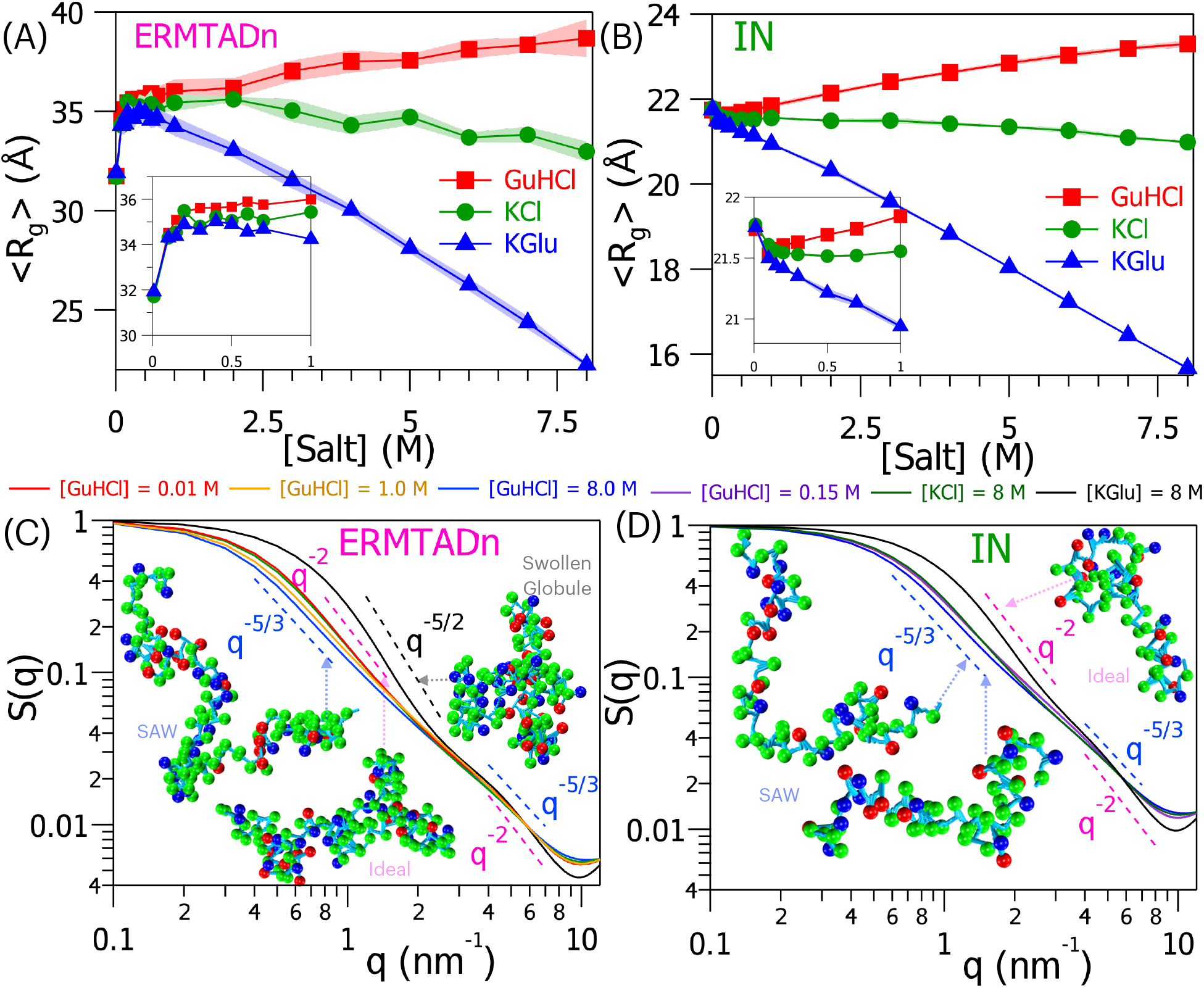
〈*R*_g_〉 of (A) ERMTADn and (B) IN computed in the presence of [*GuHCl*] (red squares), [*KCl*] (green circles) and [*KGlu*] (blue triangles). 〈*R*_g_〉 as a function of [*salt*] in low [*salt*] regime is shown in the inset. (C) *S*(*q*) of ERMTADn in the presence of [*GuHCl*] = 0.01 M (red), 1 M (yellow), 8 M (blue), [*KCl*] = 8 M (green) and [*KGlu*] = 8 M (black). (D) *S*(*q*) of IN computed in the presence of [*GuHCl*] = 0.01 M (red), 0.15 M (violet) and 8 M (blue), [*KCl*] = 8 M (green) and [*KGlu*] = 8 M (black). The dashed straight lines with *q*^−5/3^, *q*^−2^ and *q*^−5/2^ represent SAW (blue), an ideal chain (magenta) and a swollen globule (black), respectively. Representative structure(s) of IDPs are shown in bead representation. The acidic, basic and neutral residues are color coded as blue, red and green beads, respectively. of Prothymosin-*α* and Sic1 compaction in the presence of KGlu is larger compared to KCl (Fig. 3A,B).

**Figure 5:**
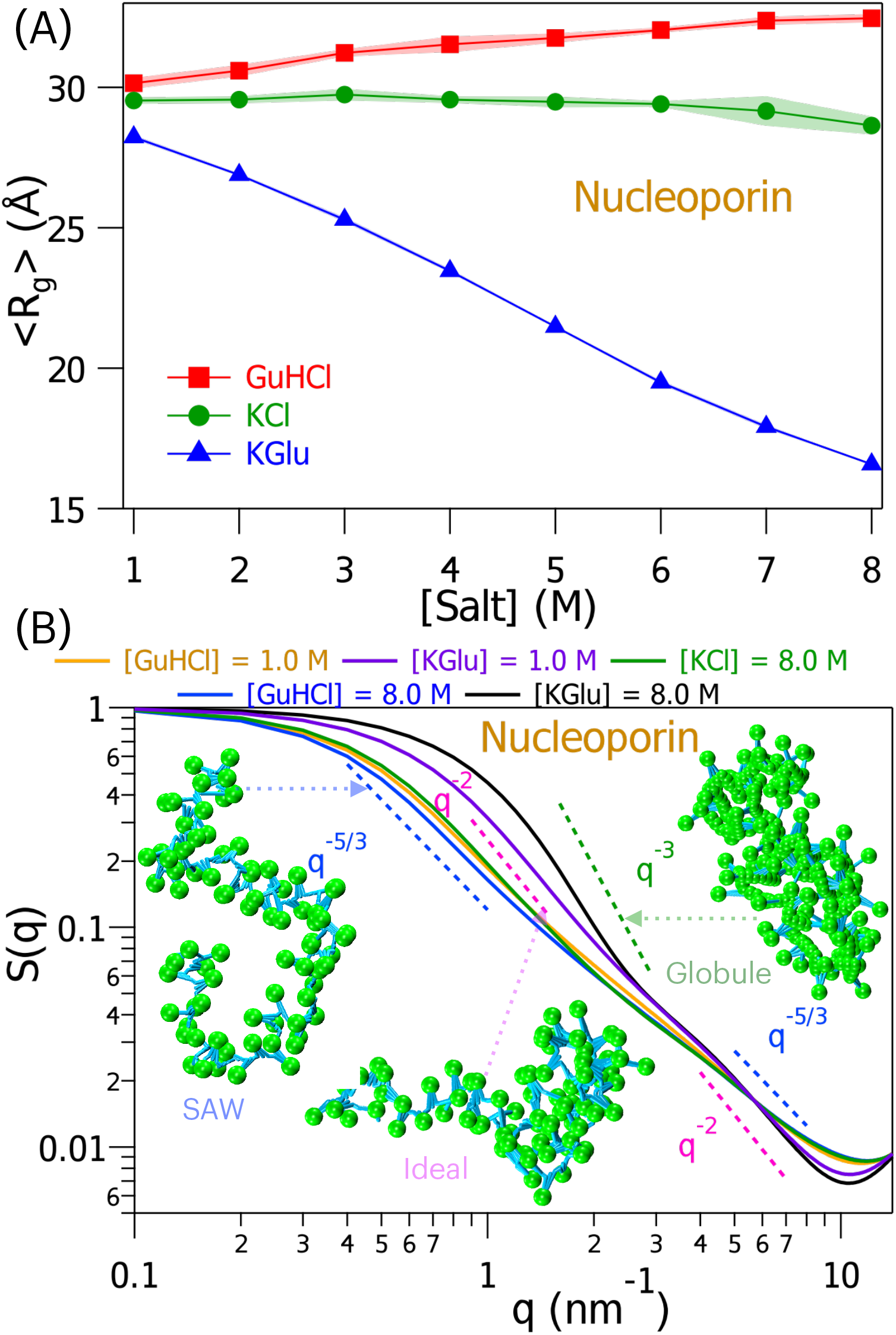
(A) 〈*R*_g_〉 of nucleoporin as a function of [*GuHCl*] (red squares), [*KCl*] (green circles) and [*KGlu*] (blue triangles). (B) *S*(*q*) of nucleoporin in the presence of [*GuHCl*] = 1 M (yellow), 8 M (blue), [*KCl*] = 8 M (green), [*KGlu*] = 1 M (violet) and 8 M (black). The dashed straight lines represent the globule (green), ideal chain (magenta) and SAW (blue) polymer chain configurations.

### Salt Induced Transitions in Polyelectrolyte-Like IDPs

We characterized the structures of polyelectrolyte-like IDPs, Prothymosin-*α* and Sic1, as a function of [*salt*]. In the low [*salt*] regime ([*salt*] < 1 M), the 〈*R*_g_〉 of the IDPs initially decreased with an increase in [*salt*] due to charge screening (Fig. 3A,B). The decrease in 〈*R*_g_〉 with an increase in [*salt*] followed the same trend obtained for polyelectrolytes in monovalent [*salt*].^34,43,44,46,85^ The decrease in 〈*R_g_*〉 for Prothymosin-*α* and Sic1 with the increase in [*salt*] from 0 to 0.7 M is ≈ 20.1 Å and 2.5 Å, respectively, and this initial compaction due to charge screening is independent for the three salts (GuHCl, KCl and KGlu) used. The effect of [*salt*] on the dimension of a charged IDP can be assessed using *FCR* or *NCPR*.^32^ Prothymosin-*α* is a strong polyelectrolyte (*FCR* ≈ 0.57, *NCPR* ≈ 0.39), whereas Sic1 is a weak polyelectrolyte (*FCR* ≈ 0.12, *NCPR* ≈ 0.12). The extent of compaction in the IDP dimensions observed for Prothymosin-*α* and Sic1 can be attributed to their strong and weak polyelectrolyte behavior.

In the high [*salt*] regime ([*salt*] > 1 M), the increase or decrease in the IDP dimensions depends on the Hofmeister effect of the salt. The salt GuHCl stabilizes expanded IDP conformations, and the dimensions of both Prothymosin-*α* and Sic1 increased with the increase in [*GuHCl*]. Whereas the salts KCl and KGlu stabilize compact IDP conformations, and the dimensions of both Prothymosin-*α* and Sic1 decreased with the increase in [*KCl*] or [*KGlu*]. KGlu strongly stabilizes compact conformations of the IDPs compared to KCl. The extent Experiments,^28^ theory^43,44,46,85^ and computer simulations^32^ on polyelectrolytes show that salt can induce structural transitions in a polyelectrolyte. In the limit [*salt*] = 0, a strong polyelectrolyte adopts a rod like structure^86^ and it can undergo a transition from rod-like to SAW in the limit of infinite [*salt*].^44^ We characterized Prothymosin-*α* and Sic1 behavior at different length scales as the [*salt*] is varied by extracting the scaling exponent *ν* from *S*(*q*) (Fig. 3C,D). In low [*salt*] (= 0.01 M), prothymosin-*α*, which is a strong polyelectrolyte behaves as a rod (*S*(*q*) ~ *q*^−1^) (Fig. 3C), whereas Sic1, which is a weak polyelectrolyte, behaves as a SAW (*S*(*q*) ~ *q*^−5/3^) (Fig. 3D). With the increase in [*salt*], Prothymosin-*α* switches from a rod-like behavior to SAW due to charge screening (Fig. 3C). In high concentration of KGlu ([*KGlu*] = 8 M), which stabilizes compact IDP conformations, Prothymosin-*α* approaches ideal polymer chain behavior (*S*(*q*) ~ *q*^−2^) (Fig. 3C), whereas Sic1 is on the verge of collapse to a globule as the value of *ν* is in between the ideal chain and collapsed globule values (1/3 < *ν* < 1/2) probably due to the finite length of Sic1 (Fig. 3D). At short length scales (*q* > 3 nm^−1^), both Prothymosin-*α* and Sic1 behave as SAWs for all [*salt*] except for [*KGlu*] = 8 M, where they exhibit ideal chain behavior (Fig. 3B,D).

We can rationalize our polyelectrolyte-like IDP results in strongly protective salt solutions using the self-consistent variational theory for the size of a polyelectrolyte chain in a poor solvent.^44^ According to this theory, the effective exclude volume, which determines the size of the polymer, depends on two terms: (1) effective two-body interaction term in the absence of charges and it is negative in poor solvents, (2) the second term is due to the charges present in the polymer, which is positive and increases with the net charge on the polymer. If both the terms balance out, then the effective exclude volume is zero, and the polymer behaves like an ideal chain. However, if the second term is larger due to a higher net charge per residue in the IDP, the effective excluded volume will be more positive, and the polymer will be swollen.

### Salt Induced Transitions in Polyampholyte-Like IDPs

A significant fraction of the IDPs are polyampholytes in nature.^87^ Based on *FCR*, ERMTADn and IN are classified as strong polyampholytes (*FCR* ≥ 0.3). The 〈*R*_g_〉 of ERMTADn increased from 31.75 Å to 36.0 Å with the increase in [*salt*] from 0 to 1 M (Fig. 4A). In contrast to ERMTADn, the 〈*R*_g_〉 of IN marginally decreased from 21.75 Å to 21.5 Å as the [*salt*] is increased to 0.15 M (Fig. 4B). The opposite trend in 〈*R*_g_〉 variation as a function of [*salt*] can be attributed to *σ*_±_ and SCD in ERMTADn (*σ*_±_ ≈ 0.007, SCD = −1.06) and IN (*σ*_±_ ≈ 0.017, SCD = 1.28) (Table S1). Expansion of ERMTADn (SCD < 0) and compaction of IN (SCD > 0) with the addition of salt in low [*salt*] are in compliance with the polyampholyte theory.^34,48,88^

ERMTADn, being a strong polyampholyte with low *σ*_±_, behaves as an ideal chain in low [*salt*] (= 0.01 M), and as the [*salt*] is increased it approaches the SAW behavior (Fig. 4C). IN is a polyampholyte with a large *σ*_±_ in low [*salt*] (= 0.01 M), and it behaves as a SAW for all concentrations of GuHCl and KCl (Fig. 4D). Although ERMTADn and IN are strong polyampholytes, the type of conformations sampled by IDPs may differ in low [*salt*] due to differences in SCD^36^ and charge patterning parameter (*κ*).^33^ ERMTADn (SCD = −1.06) is more compact than IN (SCD = 1.28) at [*salt*] = 0.01 M due to segregation of charges. The ideal chain behavior of ERMTADn (*κ* = 0.22) and SAW behavior of IN (*κ* = 0.1) at [*salt*] = 0.01 M is in accordance with the Flory random coil and excluded volume^33^ limit of random polyampholytes, respectively, based on *κ*.

In high concentration of the strong protective salt KGlu ([*KGlu*] = 8.0 M), ERMTADn undergoes compaction and it is on the verge of collapse to a globule as 1/3 < *ν* < 1/2 (Fig. 4C), and IN approaches ideal chain behavior (Fig. 4D). Both ERMTADn and IN, at short length scales (*q* > 3 nm^−1^) behave as SAW for all [*GuHCl*] and [*KCl*], and in low [*KGlu*]. For [*KGlu*] = 8.0 M, even at smaller length scales, both ERMTADn and IN exhibit ideal chain behavior.

### Salt Induced Transitions in Uncharged IDP

We investigated the effect of salts on the structure of nucleoporin, which has no charged residues. Being devoid of charged residues, modulation of coulombic interactions by salt does not apply to this IDP. The conformational ensemble of nucleoporin is affected by salt specific Hofmeister effects. The salt specific interactions with nucleoporin are similar to the other polypeptide chains investigated here. With the increase in [*GuHCl*] from 1 to 8 M, the 〈*R*_g_〉 marginally increased from ≈ 30.14 Å to ≈ 32.5 Å. In the presence of protective osmolytes, KCl and KGlu, the 〈*R*_g_〉 decreased with an increase in [*salt*]. However, the decrease in 〈*R*_g_〉 is significantly different as KCl is a weakly stabilizing agent whereas KGlu is a strongly stabilizing agent. As [*KCl*] changed from 1 to 8 M, 〈*R*_g_〉 of nucleoporin decreased by ≈ 0.9 Å. Whereas, with the increase in [*KGlu*] from 1 to 8 M, the 〈*R*_g_〉 decreased by ≈ 12.0 Å. Nucleoporin behaves like a SAW (*ν* ≈ 3/5) at all length scales for all concentrations of GuHCl, KCl and low concentration of KGlu ([*KGlu*] ¡ 1 M) as inferred from *S*(*q*). However, at higher concentration of KGlu ([*KGlu*] = 8 M) nucleoporin collapsed to compact globule like conformations (*ν* = 1/3). At smaller length scale (*q* > 4 nm^−1^), the chain exhibits ideal polymer chain behavior.

### IDP Conformations in High Concentration of Strongly Protective Salt

Depending on the polymer class to which the IDP belonged, it exhibited three kinds of behavior (*ν* = 1/2, 2/5, 1/3) in high concentrations of a strongly protective salt ([*KGlu*] = 8 M). The free energies for transferring the amino acid residues from water to a KGlu solution (*δ_g_tr__*) are positive for all the residues except for Asp and Asn.^67^ The residue Phe has the maximum positive *δ_g_tr__* value. As a result, in the KGlu solution, IDPs prefer to adopt compact conformations if they are devoid of Asp and Asn, and coil-like conformations if they are rich in Asp and Asn.

In prothymosin-*α*, there are 19 (≈ 17%) Asp, 6 (≈ 5.45%) Asn and 0 Phe residues. In IN there are 5 (≈ 8.9%) Asp, 2 (≈ 3.57%) Asn and 2 (≈ 3.57%) Phe residues. Since both Prothymosin-*α* and IN are rich in Asp and Asn, even when [*KGlu*] = 8 M they exhibit ideal chain (*ν* = 1/2) behavior (Fig. 3C and 4D). The IDP nucleoporin contains only 3 (≈ 3.7%) Asn residues, which destabilize compact conformations. All the other residues present in nucleoporin prefer compact conformations, and in addition it has 7 (≈ 8.4%) Phe residues, which strongly prefer compact conformations. As a result, when [*KGlu*] = 8 M, nucleoporin adopts a globular conformations (*ν* = 1/3) (Fig. 5B).

In Sic1 and ERMTADn, the Asn and Asp residues are more in number than the Phe residues. In Sic1, there are 6 (≈ 6.67%) Asn and 3 (≈ 3.33%) Phe residues, whereas in ERMTADn, there are 11 (≈ 9.02%) Asn, 3 (≈ 2.45%) Asp and 6 (≈ 4.92%) Phe residues. When [*KGlu*] = 8 M, both Sic1 and ERMTADn adopt neither the globule ensemble (*ν* = 1/3) nor the ideal chain ensemble (*ν* = 1/2). The value of *ν* obtained for both these IDPs is ≈ 2/5 which is in between 1/3 and 1/2 (Fig. 3D and 4C). To check if these IDPs at [*KGlu*] = 8 M are on the verge of collapse to a globule, we performed simulations in higher concentration, [*KGlu*] = 10 M, although a salt solution with this concentration is unphysical. We find that both these IDPs collapse to a globule yielding *ν* ≈ 1/3 (Fig. S4), confirming that both Sic1 and ERMTADn, at [*KGlu*] = 8 M, with *ν* ≈ 2/5 are on the verge of collapse to a globule.

When [*KGlu*] = 6 M, Sic1 and ERMTADn exhibit ideal chain behavior (*ν* ≈ 1/2). At [*KGlu*] = 8 M where the IDPs yield *ν* ≈ 2/5, we computed the probability distribution of *R*_g_, *P*(*R*_g_) to get an estimate into the population of IDPs conformations spanning between ideal chain conformations and globule conformations and compared it to those obtained at [*KGlu*] = 6 M and 10 M (Fig. S5). Computed *P*(*R*_g_) shows that at [*KGlu*] = 8 M, there is a significant IDP population in both the globule and ideal chain conformations, which can result in a *ν* value of ≈ 2/5.

### Role of Salt in the Formation of Biomolecular Condensates

Experiments^89–92^ show that salts alter the propensities of IDPs/IDRs to exhibit LLPS by modulating electrostatic and hydrophobic interactions (Fig. 6). Polymer theory based on random-phase-approximation^93^ and computer simulations^50,57,94^ demonstrated that the propensity of an IDP to exhibit LLPS is encoded in its single-molecular properties. These studies show that compaction of the IDP dimensions at the single molecular level is correlated to exhibiting LLPS. The solvent conditions (*ν* ≤ 1/2) for which the IDP dimensions are smaller than the ideal chain are conducive to observing LLPS. Hence, sequence properties such as net charge/charge patterning^94^ must play a key role in governing the LLPS propensity. The mechanism of condensate formation varies depending on [*salt*]. In low [*salt*], the LLPS propensity is controlled by electrostatic interactions independent of solvent quality. Whereas, in high [*salt*], at least ideal solvent conditions are essential for phase separation.

**Figure 6:**
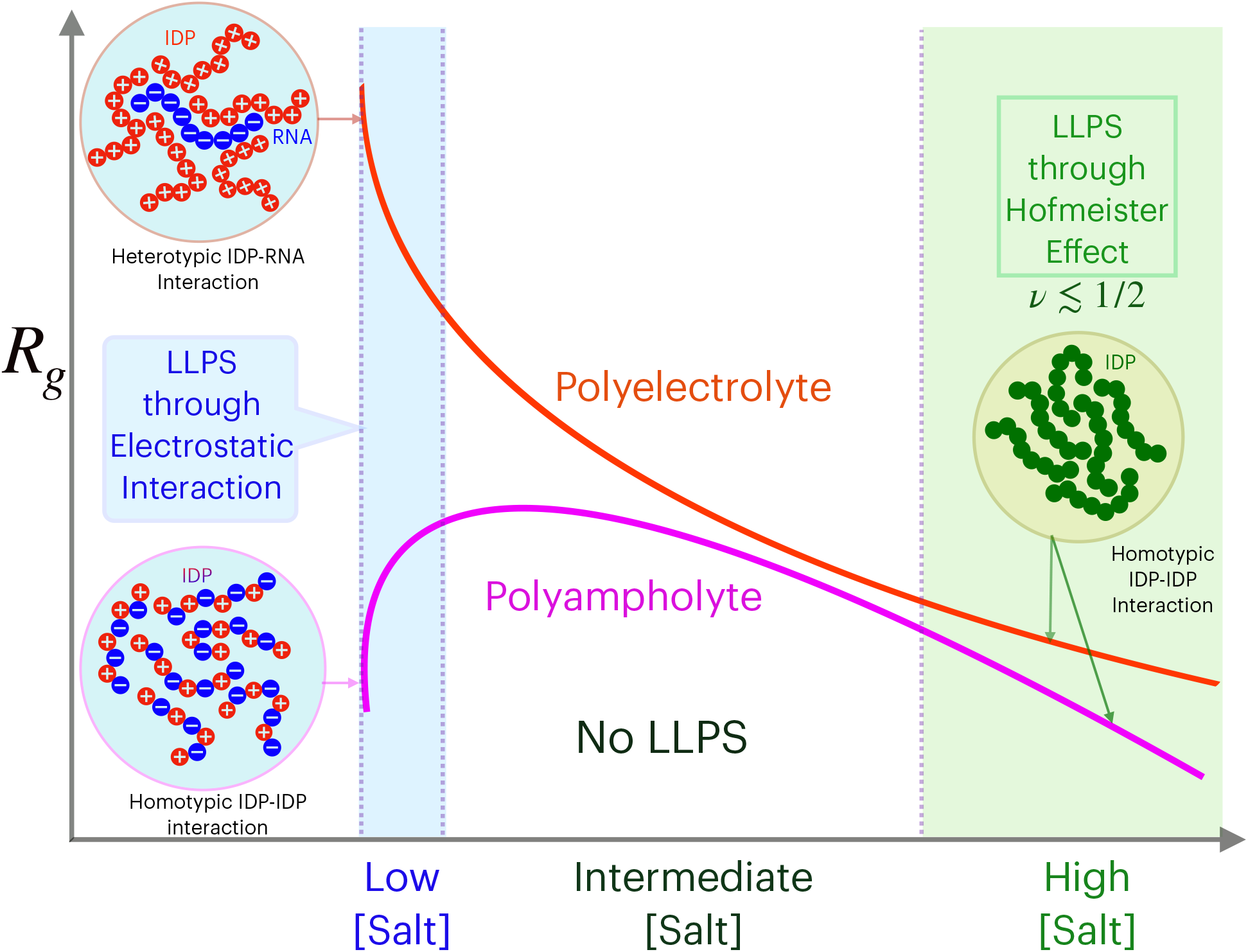
Prediction of IDP Condensate formation from single-molecule salt-dependent properties. IDPs can form condensates in low and high [*salt*]. In low [*salt*], polyelectrolyte-like IDPs exhibit LLPS in the presence of oppositely charged ligands. Whereas, polyampholytelike IDPs can undergo self-assembly without any ligands as they have both positively and negatively charged residues. In high [*salt*], the change in solvent quality due to the addition of salt leads to condensate formation by the IDPs. The quality of the solvent to the IDP modified by the addition of salt should be poorer than the ideal solvent conditions, *ν* ≤ 1/2.

In this work, we show that in the low and high [*salt*] regimes, electrostatic interactions and Hofmeister effects govern the IDP single-chain behavior, respectively. We can now predict the solvent quality of a specific salt solution to an IDP by computing the exponent *ν* from simulations performed using the IDP coarse-grained model in conditions mimicking the salt concentration. Combining this capability with the result^50,93,94^ that a correlation exists between the solvent quality to an IDP and IDPs propensity to exhibit LLPS, we can now predict whether an IDP in the semidilute regime can show LLPS when we vary the concentration of a specific salt.

In the low [*salt*] regime ([*salt*] ≲ 200 mM), polyelectrolyte-like IDPs such as prothymosin-*α* and sic1 are highly expanded due to electrostatic repulsion. Therefore, oppositely charged molecules such as RNA,^95–97^ DNA, polylysine, or polyarginine are required to induce LLPS through a network of heterotypic interactions between the IDPs and ligands.^98–100^ Conversely, polyampholyte-like IDPs such as ERMTADn and IN are stabilized due to attractive intramolecular electrostatic interactions between positively and negatively charged residues. Polyampholytes exhibit LLPS by forming a network of homotypic interactions^100^ through self-association, and ligands are not required. The IDP concentration must be in the semidilute regime to show phase separation. The critical IDP concentration required to exhibit phase separation for charged IDPs decreases with the increase in FCR, and increases with [*salt*] in low [*salt*] regime.^97^ Moreover, droplet stability of charged IDPs increases with *FCR* for a fixed [*salt*].

In the intermediate salt concentration regime (200 mM ≲ [*salt*] ≲ 2 M),^89,90^ the electrostatic interactions are screened out, and LLPS is not observed for both polyelectrolytes and polyampholyte like IDPs (Fig. 6). Uncharged IDPs like nucleoporin will not exhibit LLPS in low and intermediate [*salt*] due to the absence of charged residues and high solubility due to the polar residues present in the IDP.

IDPs can reenter the demixed phase separated state from a well-mixed state exhibiting LLPS in high concentration of stabilizing salts,^89,90^ through salt mediated Hofmeister effects^101^ (Fig. 6). Both charged and uncharged IDPs can form droplets if intramolecular monomer-monomer interaction is more favorable compared to monomer-solvent interaction at the single molecular level (*ν* ≲ 1/2).^50,93,94^ Based on the single chain properties of polyelectrolyte IDPs (Prothymosin and IN, (Fig. 3C,D)), polyampholyte IDPs (ERMTADn and sic1, (Fig. 4C,D)) and uncharged IDP (nucleoporin, (Fig. 5B)) we predict that these IDPs in the semidilute regime can reenter the phase separated state in salt solutions, [*KGlu*] ≈ 8 M, as *ν* ≲ 1/2 for these IDPs.

## Conclusion

We proposed a simulation methodology to study the effect of salts on IDPs using a reliable coarse-grained simulation model for the IDPs^49^ and experimentally measured transfer free energy of amino acids from water to the salt solutions.^25,64^ The method bridges a critical gap to understand the IDP properties in salt solutions, as salts are widely used to perturb IDP conformations to probe their role in biophysical phenomena or for use as biomaterials. Furthermore, the method overcomes the timescale problem associated with the all-atom simulations and the lack of reliable force fields to simulate high salt concentration solutions. The salt-induced transitions observed in the IDPs conformational ensemble with the variation in salt concentration depend on the polymer class of the IDP and salt identity. The role of charge composition and sequence in influencing the IDP structures in low ionic strength solutions where charge screening plays a dominant role is in accordance with the previous studies.^28,33,34^ However, in high ionic strength solutions, the transitions observed in the conformational ensembles of IDPs depend on the salt identity. Using this simulation methodology, we studied the effect of multiple salts on the single-molecule properties of IDPs, which can predict whether the salt can induce LLPS in semidilute solutions of IDPs. The model makes it feasible to directly simulate the LLPS by various IDPs in different salt solutions to understand the mechanism and dynamics of IDP condensate formation.

## Methods

To study the effect of salts on IDPs, we used a variant of the self organized polymer model for IDP (SOP-IDP).^49^ Initial structures for two bead SOP-IDP model are generated using VMD.^102^ Detailed description of the SOP-IDP energy functions are given in the Supplementary Information.

In this model, electrostatic interactions are modeled using screened Coulomb potential, and the salt specific effect on the IDP residues is taken into account using the molecular transfer model (MTM).^62,63,67^ The transfer energies of amino acids in different salts are available in Ref.^67^

We ran low friction Langevin dynamics simulations of the IDPs using the SOP-IDP model in different salt concentrations. The IDP ensemble in the presence of salt is characterized using the small-angle X-ray scattering (SAXS) intensity profiles (*I*(*q*)) and it is calculated using the equation

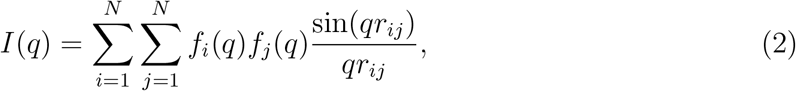

where *q* is the wave vector, *r_ij_* is the distance between the beads *i* and *j, N* is the number of beads in the IDP and *f_i_*(*q*) is the form factor of bead *i* and their values are taken from ref.^103^ The normalized structure factor,^74^ *S*(*q*), is computed using the equation

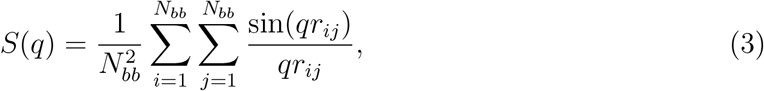

where *N_bb_* is the number of backbone beads in the IDP, and *r_ij_* is the distance between the backbone beads of residues *i* and *j*. To characterize the changes in IDP dimensions with the change in salt concentration, we computed the radius of gyration of the IDP conformations using the equation

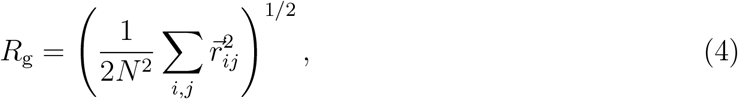

where 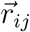 is the vector joining beads *i* and *j*.

### IDP Characterization

The IDP dimensions are characterized by the fraction of charged residues, *FCR* = (*f*_+_ + *f*_−_) and net charge per residue, *NCPR* = |*f*_+_ − *f*_−_|, where *f*_+_ and *f*_−_ are the fraction of positive and negative charged residues in the IDP. The charge asymmetry in the IDP is defined as, 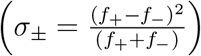. IDPs can be classified as polyelectrolytes (*FCR* > 0, *NCPR* > 0), polyampholytes (*FCR* > 0, *NCPR* ~ 0) and uncharged (*FCR* = *NCPR* = 0). Polyelectrolye and polyampholyte like IDPs are further categorized as weak (*FCR* ≤ 0.3) and strong (*FCR* > 0.3).^33,104^

In polyampholytes, segregation of positively and negatively charged residues are quantified in terms of charge patterning parameter(*κ*)^33^ and sequence charge decoration^35,36^ (SCD). *κ* is a measure of the deviation of local charge asymmetry of the blob (charged sequence segment) from the total charge asymmetry of the whole sequence. The range of *κ* is between 0 (well mixed sequence) and 1 (segregated sequence). SCD also quantifies charge patterning and it is defined as 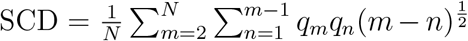, where N is the number of charged residues, *q_m_* and *q_n_* is the charge of *m*^th^ and *n*^th^ charged residue. SCD becomes more negative with increasing segregation of charged residues.

## Acknowledgement

GR acknowledges funding from Science and Engineering Research Board (EMR/2016/001356), and the National Supercomputing Mission (MeitY/R&D/HPC/2(1)/2014). HM acknowledges research fellowship from Indian Institute of Science-Bangalore. LB acknowledges research fellowship from the prime minister’s research fellows (PMRF) scheme. We acknowledge National Supercomputing Mission (NSM) for providing computing resources of “PARAM Brahma” at IISER Pune, which is implemented by C-DAC and supported by the Ministry of Electronics and Information Technology (MeitY) and Department of Science and Technology (DST), Government of India.

