## Supplementary Information for "Salt Induced Transitions in the Conformational Ensembles of Intrinsically Disordered Proteins"

### Supporting Information for "Salt Induced Transitions in the Conformational Ensembles of Intrinsically Disordered Proteins"

Hiranmay Maity,[†,‡,¶] Lipika Baidya,[†,¶] and Govardhan Reddy[*,†]

†*Solid State and Structural Chemistry Unit, Indian Institute of Science, Bengaluru, Karnataka, India 560012*

‡*Current Address: Department of Chemistry, University of Texas at Austin, Austin, TX 78712*

¶*Contributed equally to this work*

#### Materials and Methods

##### Self Organised Polymer Model for IDPs (SOP-IDPs)

We used a variant of the self-organized polymer model for intrinsically disordered proteins (SOP-IDP) to model the IDPs.[1] The SOP-IDP model is similar to the well-established self-organized polymer model with side chains (SOP-SC),[2,3] which is extensively used to study folding thermodynamics of globular proteins.[4–8] In SOP-IDP, each residue is modeled as two beads - one bead for the backbone atoms and the other bead for side chain atoms. The center of backbone bead is present at the $C_\alpha$ position, and the center of side chain bead is present at the center of mass of side chain atoms. The primary sequence of the five IDPs used in this study is shown in Table S1. To generate the initial coarse-grained coordinates ($\{\mathbf{r}\}$) for the IDP chain we used an all-atom representation of the IDP, which was constructed using the Molefacture application present in VMD.[9]

The energy function ($E_{CG}(\{\mathbf{r}\}, 0)$) of the SOP-IDP model in the absence of salt specific Hofmeister effects is the sum of bonded ($E_B$), non-bonded ($E_{NB}$) and electrostatic ($E_{ele}$) interactions. The non-bonded energy consists of local ($E_{NB}^L$) and non-local ($E_{NB}^{NL}$) interactions. The Hamiltonian of the SOP-IDP model is

$$E_{CG}(\{\mathbf{r}\}, 0) = E_B + E_{NB}^L + E_{NB}^{NL} + \lambda E_{ele}. \tag{S1}$$

If charged residues are present in the IDP, then $\lambda = 1$ else $\lambda = 0$. The bonded potential $E_B$ between two beads, which are connected by a covalent bond is modeled using the finite extensible non-linear elastic (FENE) potential,

$$E_B = -\sum_{i=1}^{N_B} \frac{k}{2} R_0^2 \log\left(1 - \frac{(r_i - r_i^0)^2}{R_0^2}\right), \tag{S2}$$

where $N_B$ is the total number of bonds in the SOP-IDP model. $r_i$ is the instantaneous bond distance between the $i^{th}$ pair of bonded beads and $r_i^0$ is their corresponding equilibrium bond distance, and $R_0$ is the maximum tolerance of bond extension/compression. The values of $r_i^0$ are set to the initial bond distance values obtained from VMD. The values of $k$ and $R_0$ are given in Table S2.

The two beads, which are not bonded by a covalent bond and separated by less than 2 residues along the polypeptide chain, interact with each other through a non-bonded local potential ($E_{NB}^L$). $E_{NB}^L$ is a purely repulsive potential accounting for excluded volume interactions to prevent unphysical overlap between the two non-bonded beads, and is given by

$$E_{NB}^L = \sum_{i=1}^{N_l} \epsilon_l \left(\frac{\sigma_i}{r_i}\right)^6, \tag{S3}$$

where $\sigma_i$ is the sum of the van der Waals (vdW) radii of $i^{th}$ pair of non-bonded beads and $\epsilon_l$ is the strength of repulsive interaction. The value of $\epsilon_l$ is given in Table S2 and values of vdW radii for each amino acid residue are listed in Table S3.

The beads, which are separated by more than two residues, interact through non-bonded non-local interaction potential, $E_{NB}^{NL}$, which is modeled using

$$\begin{aligned} E_{NB}^{NL} = & \sum_{i=1}^{N_{bb}} \omega \times 300 k_{\mathrm{B}} \times \mid 0.7 - \epsilon_i^{bb} \mid \left[ \left( \frac{\sigma^{bb}}{r_i} \right)^{12} - 2 \left( \frac{\sigma^{bb}}{r_i} \right)^{6} \right] \\ & + \sum_{i=1}^{N_{bs}} \omega \times 300 k_{\mathrm{B}} \times \mid 0.7 - \epsilon_i^{bs} \mid \left[ \left( \frac{\sigma_i^{bs}}{r_i} \right)^{12} - 2 \left( \frac{\sigma_i^{bs}}{r_i} \right)^{6} \right] \\ & + \sum_{i=1}^{N_{ss}} \omega \times 300 k_{\mathrm{B}} \times \mid 0.7 - \epsilon_i^{ss} \mid \left[ \left( \frac{\sigma_i^{ss}}{r_i} \right)^{12} - 2 \left( \frac{\sigma_i^{ss}}{r_i} \right)^{6} \right]. \end{aligned} \quad \text{(S4)}$$

The first, second and third terms of Eq. S4 correspond to the backbone-backbone, backbone-side chain and side chain-side chain interactions energies, respectively. $N_{bb}$, $N_{bs}$ and $N_{ss}$ denote the number of interaction pairs present between backbone-backbone, backbone-side chain and side chain-side chain beads, respectively. $r_i$ is the distance between $i^{th}$ pair of beads and $k_{\mathrm{B}}$ is the Boltzmann constant. $\sigma^{bb}$ is the diameter of the backbone bead, which is taken as 3.8 Å. $\sigma_i^{bs}$ and $\sigma_i^{ss}$ are the sum of bead radii for the $i^{th}$ pair of backbone-side chain and side chain-side chain beads, respectively. $\sigma_i^{bs}$ and $\sigma_i^{ss}$ are computed using the bead radii listed in Table S3. $\epsilon_i^{bb}$, $\epsilon_i^{bs}$ and $\epsilon_i^{ss}$ are the strength of backbone - backbone, backbone - side chain and side chain - side chain interactions, respectively. We used Betancourt - Thirumalai statistical potential for $\epsilon_i^{ss}$, which was initially proposed for the side chain - side chain interactions of globular proteins.[10] However, for IDPs we used a rescaling factor $\omega$ ($0 < \omega < 1$) to weaken the strength of interactions. The value of $\epsilon_i^{bb}$ is approximated to be the Betancourt - Thirumalai statistical potential corresponding to the interaction between two Gly residues. $\epsilon_i^{bs}$ is approximated as the statistical potential for the interaction between Gly and the side chain corresponding to the other residue. We have used an $\omega$ value for which the experimental and simulated SAXS profiles of the IDPs are in agreement. The value of $\omega$ used for all the IDPs is given in Table S2. We have used the truncated and shifted form of LJ potential using a cutoff distance 30 Å beyond which the interaction between the two beads is neglected.

If an IDP contains charged residues, the beads corresponding to the side chains of the charged residues interact through a screened Coulomb potential given by

$$E_{ele} = \sum_{i=1}^{N_c-1} \sum_{j>i}^{N_c} \frac{q_i q_j \exp(-\kappa r_{ij})}{\epsilon r_{ij}}, \tag{S5}$$

where $N_c$ is the total number of charged residues present in the IDP, $r_{ij}$ is the distance between charged beads $i$ and $j$, $q_i$ and $q_j$ are the point charges measured in units of electron charge placed on the centers of the side chain beads $i$ and $j$, respectively. At neutral pH, $q_i$ is $+1$ for positively charged Lys and Arg residues and -1 for negatively charged Asp and Glu residues. The inverse Debye length, $\kappa$, is computed for 150 mM monovalent salt concentration. We have used the dielectric constant of the medium $\epsilon = 78.0\epsilon_0$, where $\epsilon_0$ $(= 1.0)$ is the permittivity of vacuum.

#### Molecular Transfer Model (MTM)

Salts in low concentration affect the IDP conformations by screening the electrostatic interactions and this effect is independent of the salt identity but depends on the salt concentration ([salt]). Whereas in high concentration, salts affect the IDP conformations through salt specific Hofmeister effects. The non-specific Coulombic interaction is modeled by the $E_{ele}$ potential (Eq. S5).

To introduce salt specific interactions we used molecular transfer model (MTM), which was extensively used previously to study folding thermodynamics of globular proteins.[11] In MTM, the modified energy function of an IDP conformation with coordinates ($\{\mathbf{r}\}$) and salt concentration $[salt]$ is given by

$$E_{CG}(\{\mathbf{r}\},[salt]) = E_{CG}(\{\mathbf{r}\},0) + \Delta G_{tr}(\{\mathbf{r}\},[salt]), \tag{S6}$$

where $\Delta G_{tr}(\{\mathbf{r}\},[salt])$ is the transfer free energy associated with the transferring of an IDP conformation from water to a salt solution with concentration $[salt]$, and is given by

$$\Delta G_{tr}(\{\mathbf{r}\},[salt]) = \sum_{i=1}^{N_{res}} \delta g_{tr}^{bb}([salt]) \frac{\alpha_i^{bb}(\{\mathbf{r}\})}{\alpha_{Gly-i-Gly}^{bb}} + \sum_{i=1}^{N_{res}} \delta g_{tr,i}^{sc}([salt]) \frac{\alpha_i^{sc}(\{\mathbf{r}\})}{\alpha_{Gly-i-Gly}^{sc}}, \tag{S7}$$

where $N_{res}$ is the total number of residues in the IDP, $\delta g_{tr}^{bb}([salt])$ and $\delta g_{tr,i}^{sc}([salt])$ are the transfer free energies of backbone bead and side chain bead of $i^{th}$ residue from water to a salt solution $[salt]$, respectively, $\alpha_i^{bb}(\{\mathbf{r}\})$ and $\alpha_i^{sc}(\{\mathbf{r}\})$ are the solvent accessible surface area (SASA) of the backbone and side chain bead of residue $i$ in the protein chain, $\alpha_{Gly-i-Gly}^{bb}$ and $\alpha_{Gly-i-Gly}^{sc}$ are the SASA of the backbone and side chain beads of the same amino acid residue $i$ in the tripeptide $Gly-i-Gly$. The values of $\delta g_{tr}^{bb}([salt])$, $\delta g_{tr,i}^{sc}([salt])$, $\alpha_{Gly-i-Gly}^{bb}$ and $\alpha_{Gly-i-Gly}^{sc}$ are available in ref.[11] The SASA of IDP conformations is calculated using the method described by Wodak and Janin.[12]

#### Simulation Details and Data Analysis

We carried out low friction Langevin dynamics[13] simulation at temperature, $T = 300$ K in presence of salts to compute the average thermodynamic properties of IDPs. The equation motion in Langevin dynamics is given by,

$$m\ddot{\vec{r}} = -\zeta\dot{\vec{r}} + \vec{F}_C + \vec{\Gamma}, \tag{S8}$$

where $m$ is the mass of protein beads, $\zeta$ is the friction coefficient of the solvent medium, $\vec{F}_C$ is the deterministic force given by $-\frac{\partial E_{CG}(\{\mathbf{r}\},[salt])}{\partial \vec{r}_i}$, and $\vec{\Gamma}$ is the random force with Gaussian noise characterized by $\left\langle \vec{\Gamma}(t) \cdot \vec{\Gamma}(t+nh) \right\rangle = \frac{2\zeta k_B T}{h}\delta_{0,n}$ where $n = 0, 1, \cdots$, $\delta_{0,n}$ is the Kronecker delta function and $k_B$ is the Boltzmann constant. We integrated Eq. S8 using the velocity Verlet algorithm. We used $\zeta = 0.05\ m/\tau_{LD}$ and an integration timestep, $h = 0.005$ $\tau_{LD}$, where, $\tau_{LD}$ $\left(= \sqrt{\frac{m_0 a_0^2}{\epsilon_h}}\right)$ is the unit of time used to advance the Langevin dynamics simulations. The average mass of each bead ($m_0$), characteristic unit of length ($a_0$) and energy ($\epsilon_h$) are taken as 1.8 $\times 10^{-22}$ g, 1 Å and 1 kcal/mol. The value of $\tau_{LD}$ in real time unit is $\approx$ 1.3 ps. The length of the simulations time for each IDP at a particular salt concentration is at least $3 \times 10^6$ $\tau_{LD}$. By computing the autocorrelation function of square of end-to-end distance, we show that the simulation time is adequate to sample the relevant conformational space of IDPs (Fig. S6). The initial $10^3$ $\tau_{LD}$ steps are discarded and not used to compute thermodynamic properties.

To test whether the simulation time is adequate to study IDP thermodynamic properties, we computed the autocorrelation function of square of end-to-end distance, $R_{ee}^2$, defined as

$$A_{R_{ee}^2}(t) = \frac{\langle R_{ee}^2(t_0) R_{ee}^2(t_0 + t)\rangle - \langle R_{ee}^2\rangle^2}{\langle R_{ee}^4\rangle - \langle R_{ee}^2\rangle^2}, \quad \text{(S9)}$$

where $R_{ee}^2(t_0)$ and $R_{ee}^2(t_0+t)$ are the squared end-to-end distance at time $t_0$ and $t_0+t$, $\langle ... \rangle$ denotes the ensemble average, $\langle R_{ee}^2\rangle$ and $\langle R_{ee}^4\rangle$ are the $2^{nd}$ and $4^{th}$ moments of $R_{ee}$, respectively.

The simulated small-angle X-ray scattering (SAXS) intensity ($I(q)$) profiles for the IDPs are calculated using the expression

$$I(q) = \sum_{i=1}^{N}\sum_{j=1}^{N} f_i(q) f_j(q) \frac{\sin(q r_{ij})}{q r_{ij}}, \quad \text{(S10)}$$

where $N$ is the total number of beads, $q$ is the scattered wave vector, $f_i(q)$ is the form factor of bead $i$, and $r_{ij}$ is the distance between the beads $i$ and $j$. The values of $f_i(q)$'s are taken from ref.[14] We evaluated the normalized structure factor, $S(q)$, using the backbone beads and it is given by

$$S(q) = \frac{1}{N_{bb}^2} \sum_{i=1}^{N_{bb}} \sum_{j=1}^{N_{bb}} \frac{\sin(qr_{ij})}{qr_{ij}}, \tag{S11}$$

where $N_{bb}$ is the total number of backbone beads and $r_{ij}$ is the distance between the beads $i$ and $j$.

The radius of gyration, $R_g$, of the IDPs is given by

$$R_g = \left( \frac{1}{2N^2} \sum_{i,j} \vec{r}_{ij}^{\,2} \right)^{1/2}, \tag{S12}$$

where $\vec{r}_{ij}$ is the vector joining beads $i$ and $j$ and $N$ is the number of beads in the IDP.

Table S1: Primary sequences of nucleoporin, sic1, ERMTADn, IN, and prothymosin-$\alpha$ are listed from top to bottom, respectively. $f_+$ and $f_-$ are the fraction of positive and negatively charged residues. The positively charged residues Lys (K) and Arg (R) are given in red, whereas the negatively charged residues Asp (D) and Glu (E) are given in blue. FCR ($f_+ + f_-$) and NCPR ($|f_+ - f_-|$) are the fraction of charged residues and net charge per residue, respectively. Charge asymmetry ($\sigma_\pm$) is defined as $\frac{(f_+ - f_-)^2}{(f_+ + f_-)}$. Nucleoporin is electrostatically neutral IDP with zero charged residue. Sic1 and prothymosin-$\alpha$ are polyelectrolytes, whereas ERMTADn and IN are polyampholytes.

| Protein | Sequence | $N_{res}$ | $f_+$ | $f_-$ | FCR | NCPR | $\sigma_\pm$ |
|---|---|---|---|---|---|---|---|
| Nucleoporin | GCPSASPAFGANQTPTFGQSQ<br>GASQPNPPGFGSISSSTALFPT<br>GSQPAPPTFGTVSSSSQPPV<br>FGQQPSQSAFG SGTTPNA | 81 | 0.0 | 0.0 | 0.0 | 0.0 | 0.0 |
| Sic1 | MTPSTPPRSRGTRYLAQPSGN<br>TSSSALMQGQKTPQKPSQNL<br>VPVTPSTTKSFKNAPLLAPP<br>N S NMGMTSPFNGLTSPQRSP<br>FPKSSVKRT | 90 | 0.12 | 0.0 | 0.12 | 0.12 | 0.122 |
| ERMTADn | MDGFYDQQVPFMVPGKSRSE<br>ECRGRPVIDRKRKFLDTDLAH<br>DSEELFQDLSQLQEAWLAEAQ<br>VPDDEQFVPDFQSDNLVLAPP<br>PTKIKRELHSPSSELSSCSHEQ<br>ALGANYGEKCLYNY CA | 122 | 0.13 | 0.18 | 0.31 | 0.05 | 0.007 |
| IN | FLDGIDKAQEEHEKYHSNWR<br>AMASDFNLPPVVAKEIV<br>ASCDKCQLKGEAMHGQVDC | 56 | 0.11 | 0.18 | 0.29 | 0.07 | 0.017 |
| Prothymosin-$\alpha$ | MSDAAVDTSSEITTKDLKEKK<br>EVVEEAENGRDAPANGNANE<br>EENGEQEADNEVDEEEEEGG<br>EEEEEEEEGDGEEEDGDEDEE<br>AESATGKRAAEDDEDDDVDT<br>KKQK TDEDD | 110 | 0.09 | 0.48 | 0.57 | 0.39 | 0.27 |

Table S2: Parameters used in SOP-IDP Model

| Parameter | Value |
|---|---|
| $k$ | 20.0 kcal/(mol.Å$^2$) |
| $R_0$ | 2.0 Å |
| $\epsilon_l$ | 1.0 kcal/mol |
| $\epsilon_{bb}$ | -0.2 |
| $\omega$ | 0.12 |
| $\epsilon$ | 78 |

Table S3: Backbone and side chain bead radii of amino acid residue

| Bead | vdW radius (Å) |
|---|---|
| backbone | 1.90 |
| Gly | 0.5 |
| Ala | 2.52 |
| Val | 2.93 |
| Leu | 3.09 |
| Ile | 3.09 |
| Met | 3.09 |
| Phe | 3.18 |
| Pro | 2.78 |
| Ser | 2.59 |
| Thr | 2.81 |
| Asn | 2.84 |
| Gln | 3.01 |
| Tyr | 3.23 |
| Trp | 3.39 |
| Asp | 2.79 |
| Glu | 2.96 |
| Hsea | 3.04 |
| Hsd | 3.04 |
| Lys | 3.18 |
| Arg | 3.28 |
| Cys | 2.74 |

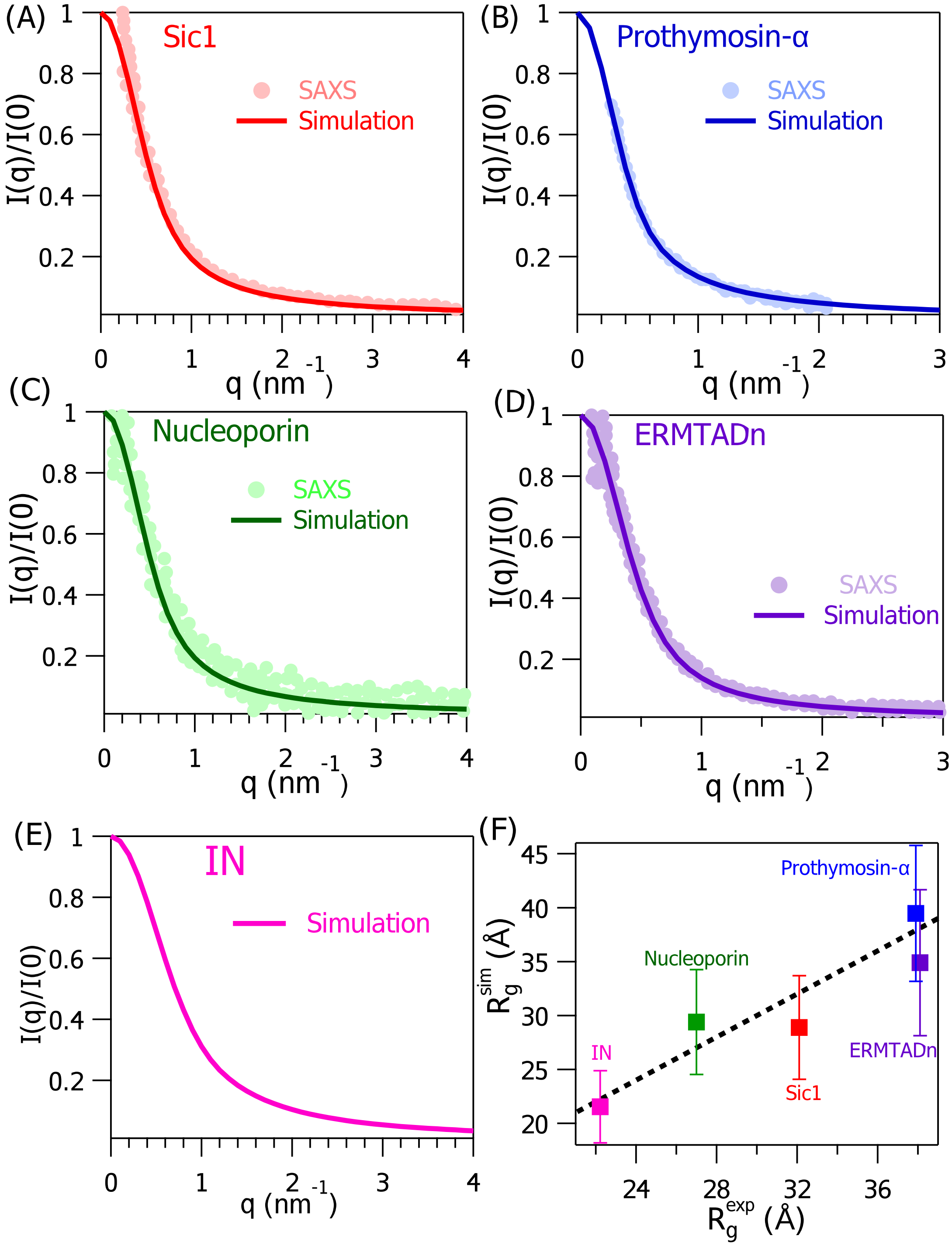


Figure S1: Normalized scattering intensity $(I(q)/I(0))$ of scattered wave vector, $q$ from experiments and simulations are plotted for (A) sic1, (B) prothymosin-$\alpha$, (C) nucleoporin, (D) ERMTADn and (E) IN. Experimental data is not available for IN. (F) Average radius of gyration of IDPs from simulation, $R_g^{sim}$ are plotted against experimentally measured values, $R_g^{exp}$ . The Pearson correlation coefficient between $R_g^{exp}$ and $R_g^{sim}$ is $\approx 0.93$.

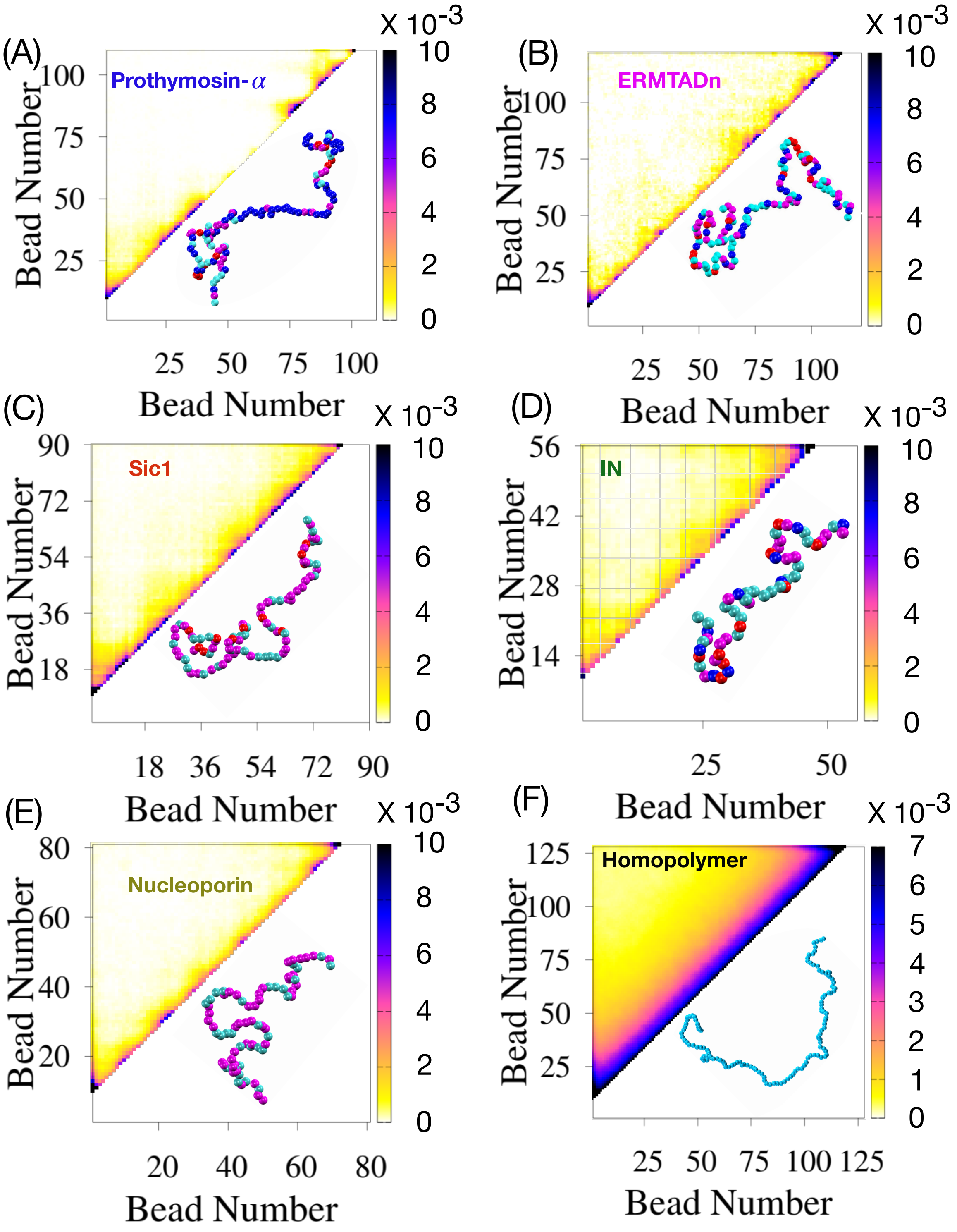


Figure S2: Probability of nonlocal contact formation between two residues which are separated by more than 8 residues are plotted at temperature, T = 300 K and [salt] = 0.15 M for (A) Prothymosin-$\alpha$, (B) ERMTADn, (C) sic1, (D) IN and (E) nucleoporin. The contact frequency obtained for all pairs of residues for IDPs is highly heterogeneous for a given sequence. The color code of beads - blue (acidic), red (basic), cyan (nonpolar) and magenta (polar) denote nature of the residues. Probability of nonlocal contact formation for homopolymer in (F).

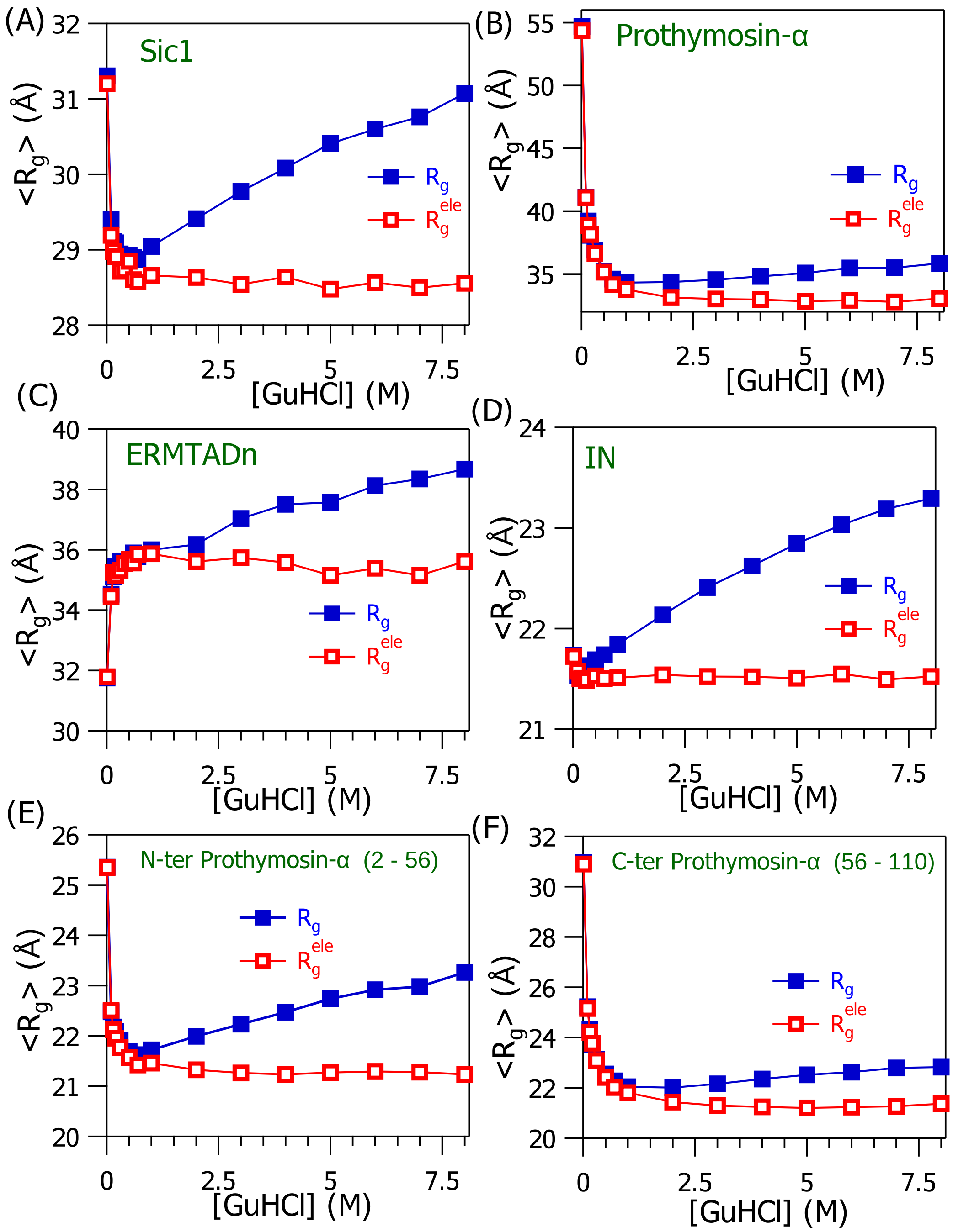


Figure S3: Dissecting the electrostatic charge screening and Hofmeister effects of GuHCl on the IDPs. The average radius of gyration, $\langle R_g^{ele} \rangle$ and $\langle R_g \rangle$ are computed using the energy functions Eq. S1 and S6, respectively. The data for $\langle R_g^{ele} \rangle$ and $\langle R_g \rangle$ are shown in open red squares and solid blue squares, respectively. The data is plotted for (A) sic1 (B) prothymosin-$\alpha$ (C) ERMTADn (D) IN (E) N-terminal segment of prothymosin-$\alpha$ (residue 2 to 56) and (F) C-terminal segment of prothymosin-$\alpha$ (residue 56 to 110). Initially, $\langle R_g^{ele} \rangle$ rapidly decreases or increases as [GuHCl] increases from 10 mM to 1 M, and for [GuHCl] $>$ 1 M it remains almost constant. When the transfer energy is included in the energy function to take into account the Hofmeister effects, $\langle R_g \rangle$ increases gradually as the [GuHCl] increases from 1 M to 8 M. In the high [salt] regime, salt induced structural changes occur through Hofmeister effects.

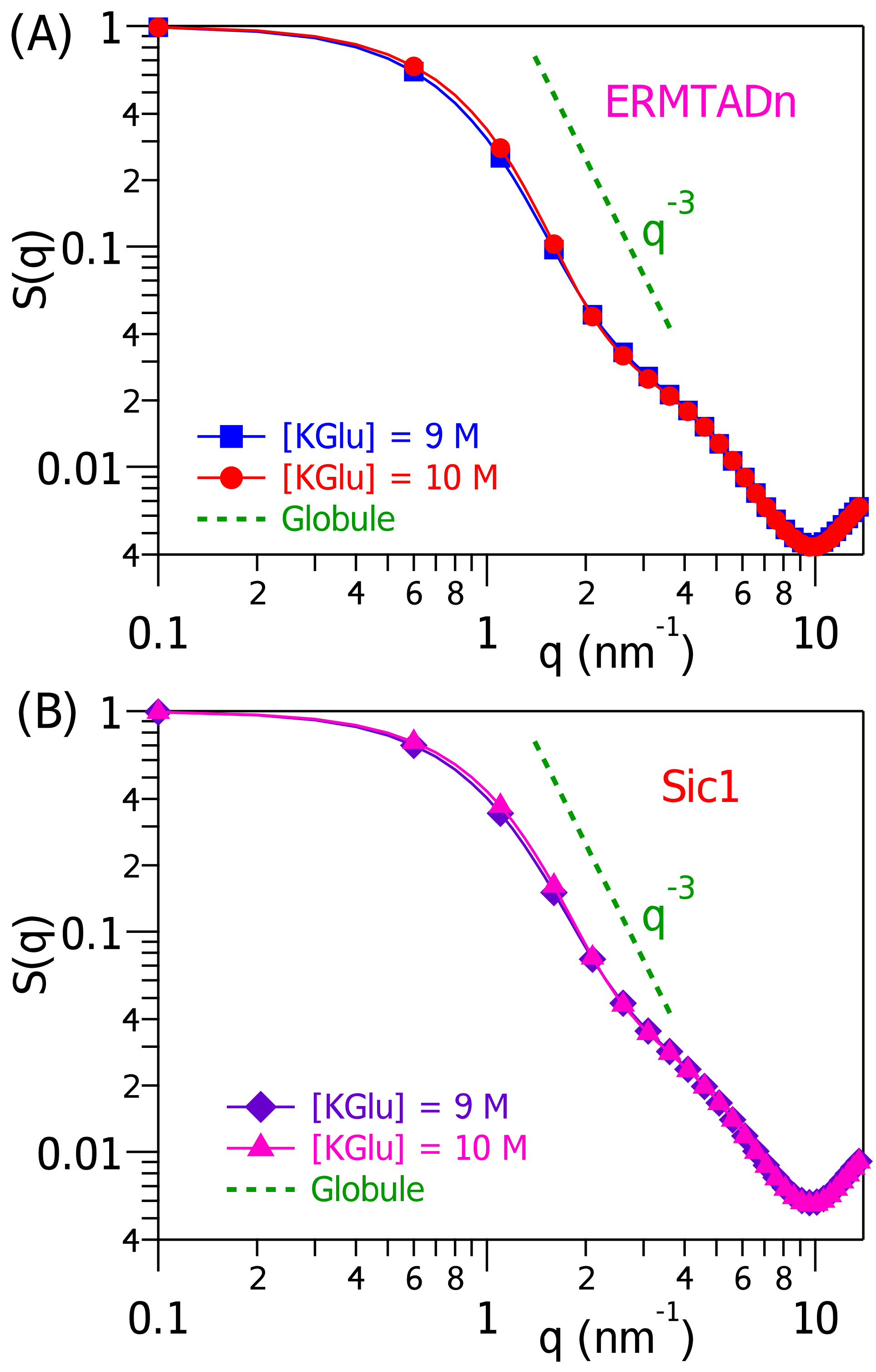


Figure S4: Structure factor, $S(q)$ as a function of wave vector, $q$ in $[KGlu] = 9$ M and 10 M for ERMTADn in (A) and sic1 in (B) scale with $q^{-3}$ which denote the dimension of IDPs are globule (green dotted line).

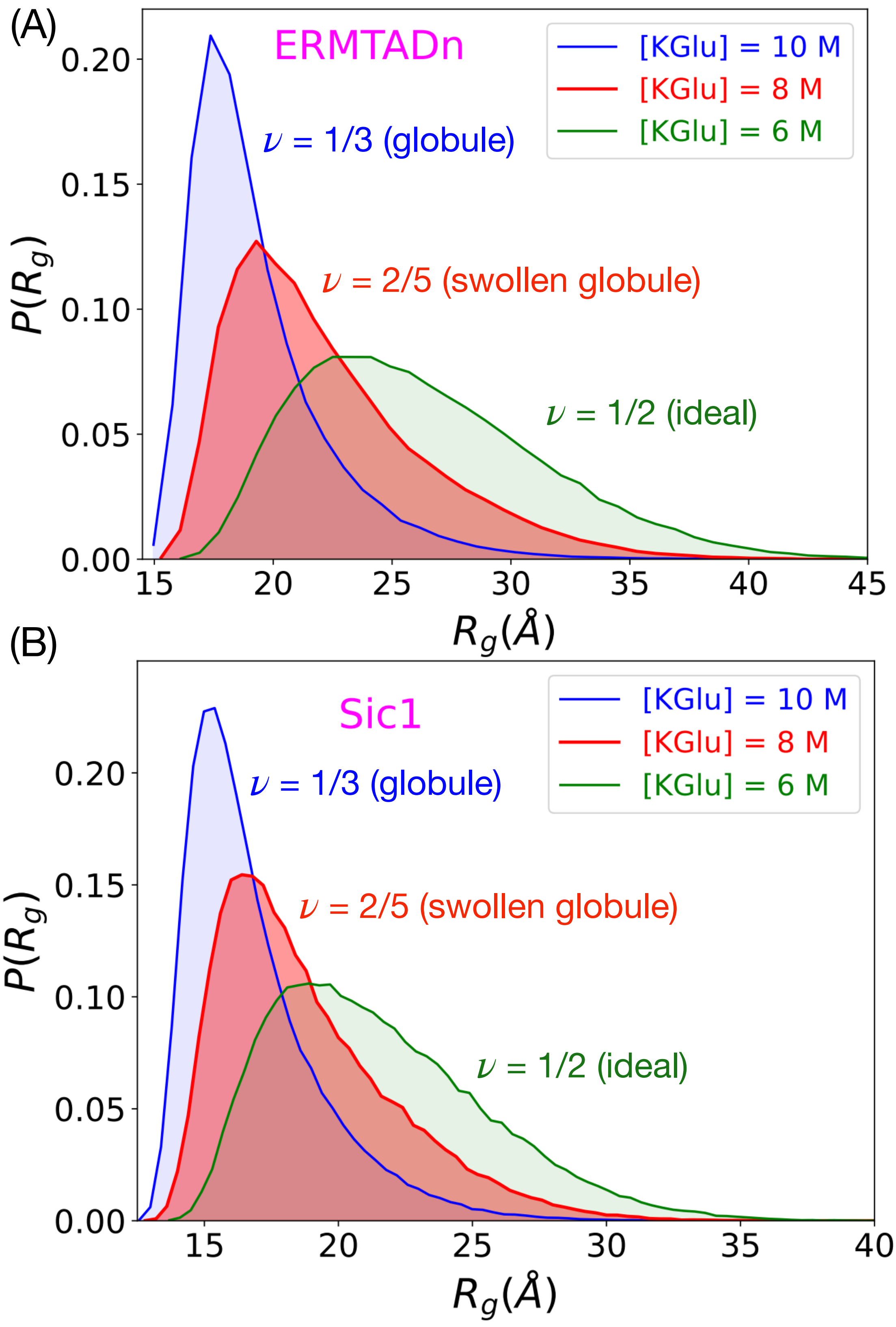


Figure S5: Probability distribution of radius of gyration, $R_g$ in $[KGlu] = 6$, 8, 10 M for ERMTADn in (A) and sic1 in (B).

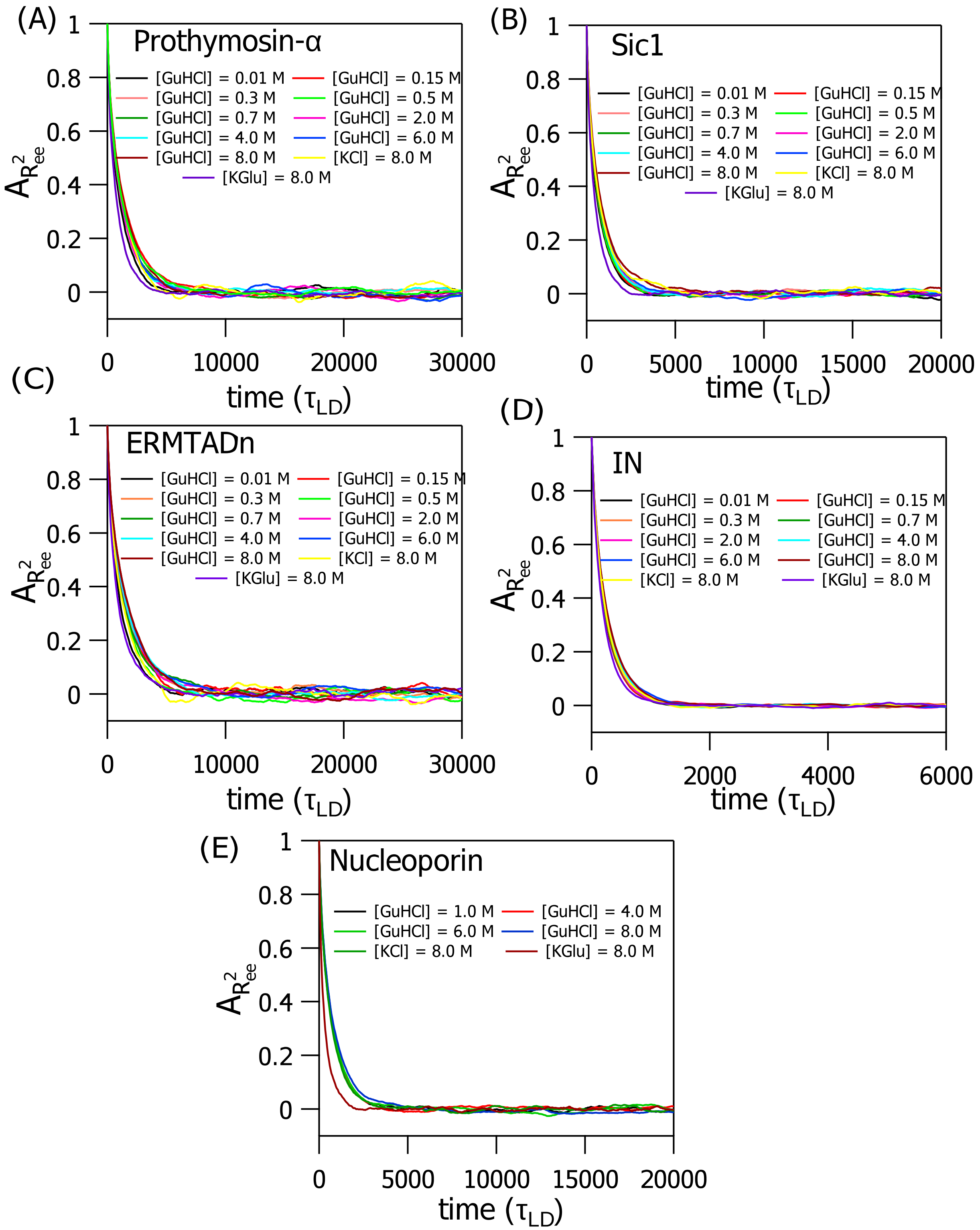


Figure S6: End-to-end autocorrelation function, $A_{R^2_{ee}}$ as a function of time, $\tau_{\rm LD}$ is plotted for (A) prothymosin-$\alpha$ (B) sic1 (C) ERMTADn (D) IN and (E) nucleoporin. The lines with different colors correspond to different $[salt]$. The autocorrelation of end-to-end distance decays $< 10^3\tau_{\rm LD}$. Data from the initial $10^3\tau_{\rm LD}$ time steps are discarded and not used for analysis. We used at least $3 \times 10^6\tau_{\rm LD}$ time steps of simulation data for analysis.
